# Aquavert – Imaging and Microfluidics for Vertical Swimming of Microorganisms

**DOI:** 10.1101/2024.09.07.611807

**Authors:** Haley B. Obenshain, Isaias Zarate, Olivia Hedman-Manzano, Jared Goderich, Sungho Lee, Bryant A. Lopez, Emma Varela, Ga-Young Kelly Suh, Douglas A. Pace, Siavash Ahrar

**Author notes:** **Corresponding Authors Contacts** Siavash Ahrar (Ph.D.), Mail: Department of Biomedical Engineering, California State University Long Beach, Long Beach, CA 90840, United States of America., Douglas A. Pace (Ph.D.), Mail: Department of Biological Sciences, California State University Long Beach, Long Beach, CA 90840, United States of America. Citation*: Obenshain, et al., Aquavert – Imaging and Microfluidics for Vertical Swimming of Microorganisms, DOI:000000/11111.

## Abstract

Investigating aquatic microorganisms’ swimming and feeding behaviors under well-controlled conditions is of great interest across multiple disciplines. Thus, broader access to resources that enable these investigations is desirable. Given the organisms’ microscopic dimensions, an ideal system should combine microscopy to visualize and fluidics to control and modulate their environments. We report an integrated device (Aquavert) that combines DIY microscopy and microfluidics for biomechanical investigations of marine microorganisms, emphasizing vertical swimming. The DIY microscope was developed for modularity, and imaging chambers were secured in vertical orientations (either in portrait or landscape mode). Fluid channels were used to introduce flow and fluid segmentation while remaining upright. Fluid segmentation established two distinct environments (e.g., with and without algae) in neighboring regions inside a chamber. System application with multiple marine larvae (sand dollars, sea urchins, and starfish) and introduction of unicellular algae were demonstrated. Finally, the device’s capabilities were extended to fluorescence imaging to visualize tracer beads. The role of gravity is often ignored in conventional plate or microfluidic experiments. Beyond the current application, Aquavert enables investigations of the behavior and physiology of microorganisms where the role of gravity is critical.

## 1 Introduction

Broad access to microscopy and imaging tools is desirable across biological investigations. The Do-It-Yourself (DIY) movement has provided frameworks and resources to implement inexpensive, robust, and customizable microscopes [1, 2, 3]. Unlike conventional microscopes, recent DIY systems have sought to fulfill unique requirements optimized for a particular need. For example, devices for tracking organisms (Trackoscope) [4], examining biodiversity of insects (Entomoscope) [5], investigating microplastic pollution (EnderScope) [6], broadening access to citizen oceanography (Planktoscope)[7], providing orthogonal views of organisms (GLUBscope) [8], and 3D mesoscopic imaging of fixed and live samples in the mm–cm scale (OPTImAL) [9] have been demonstrated. Additional efforts led by Prakash lab have significantly advanced DIY microscopy across multiple applications and requirements [10, 11, 12]. We describe an integrated device that combines DIY microscopy and vertically oriented channels to investigate microorganisms’ swimming and feeding behaviors.

Investigating the swimming and feeding behaviors of marine microorganisms is of great interest. However, field investigations of these behaviors are challenging. Therefore, laboratory investigations and access to corresponding resources are needed. In one approach, laboratory investigations use particle image velocimetry (PIV) to visualize organisms’ swimming and the corresponding flow profile. For example, Wheeler et al. used tanks (90 cm× 45 cm × 45 cm), an IR laser light source, and tracer particles to visualize the swimming behavior of marine larvae [13, 14]. In another example using the PIV approach, Wong et al. used a custom made glass chamber (25 mm×75 mm×5 mm) and an array of white LEDs to investigate the swimming kinematics of barnacle naupliar larvae [15]. In another approach, Durham and co-authors developed an elegant system to investigate the gyrotactic trapping of phytoplankton during vertical movement [16]. Using a belt-driven mechanism, investigators created a vertical gradient in a horizontal velocity profile inside a chamber (36.7 cm×21.6 cm×1 cm). More recently, Krishnamurthy and co-authors developed a circular chamber integrated with microscopy to investigate the swimming behavior of various organisms at an ecological scale [17, 18]. Our investigation considered the following requirements as gaps in the existing tools. First, while large chambers provide ample swimming space, their maintenance, operation, and compatibility with microscopy are complex. Thus, miniaturization that enables vertical swimming is desirable. Second, the ability to provide well-controlled external flows or fluid segmentation (i.e., regions that have different characteristics such as food concentrations, pH, temperature, salinity, and toxins) has remained absent. Third, the ability to customize and exchange components was considered, given the diversity of size and morphologies of marine organisms. An integrated device that combines DIY microscopy for vertically oriented microfluidics was developed to achieve these requirements.

We provide instructions for reproducing the integrated device (Aquavert) and proof-of-use experiments. The device is divided into two modules: imaging and fluidics. The imaging system was positioned to obtain data from vertically oriented microfluidics. We prioritized the modularity and customizability of optical components. For example, depending on the experiment, different objectives were used (2X, 4X, and 10X presented), or additional optical filters were introduced. Both the application of a laptop and a Raspberry Pi were demonstrated to control the camera. Finally, fluorescence imaging with the system was demonstrated. Multiple microfluidic designs were developed for well-controlled flow and fluid segmentation. The length scales of channels were longer (4-10 mm wide and 65 mm long) than typical microfluidics to accommodate marine larvae and minimize concerns related to confinement. Given the vertical swimming behaviors of marine organisms, two unique chamber holders were developed to secure the vertical chambers in portrait or landscape orientations. Aquavert was used to investigate the vertical swimming behaviors with and without external flow in these orientations in multiple marine larvae, including those of *Dendraster excentricus* (eccentric sand dollar) and *Pycnopodia helianthoides* (sunflower sea star), both with and without external flow, across different orientations and chip designs.

## 2 Results

### 2.1 Aquavert Concept

Aquatic zooplankton display a diversity of morphologies. Aquavert provides an approach to study the functional consequences of these morphologies. In particular, vertical swimming is of great interest [19, 20]. Previous investigations usually use large chambers that are not readily compatible with microscopy. Figure 1 summarizes the system-level diagram and images of the Aquavert device and highlights the orientation of chamber holders. We sought to use standard microfluidics instead of custom made chambers, due to their broad availability across multiple disciplines. The chambers were then positioned vertically, given the organisms’ swimming behaviors. It is essential that the chambers have well-controlled fluid flows allowing for flow segmentation. System modularity, which enables various experiments and organisms (particularly given various marine organisms’ morphologies), was considered the primary design principle. Ease of replication, size reduction, and cost minimization were also considered. In our experience, access to imaging, not fluidics, is the bottleneck for scaling microfluidic investigations. Therefore, improving access to imaging through cost reduction could enable parallel research activities. In the context of marine larvae, this parallelization is also valuable since considering the age of the larvae (days post fertilization) is an essential factor in determining the morphology, stage, and corresponding behaviors. We combined off-the-shelf components and open-sourced DIY parts to achieve these design goals. Moreover, the application of a Raspberry Pi controller instead of a computer was demonstrated. The report provides instructions for building the Aquavert device and proof-of-use application experiments. To simplify the instructions, we have divided the device into imaging and fluidics modules.

**Figure 1:**
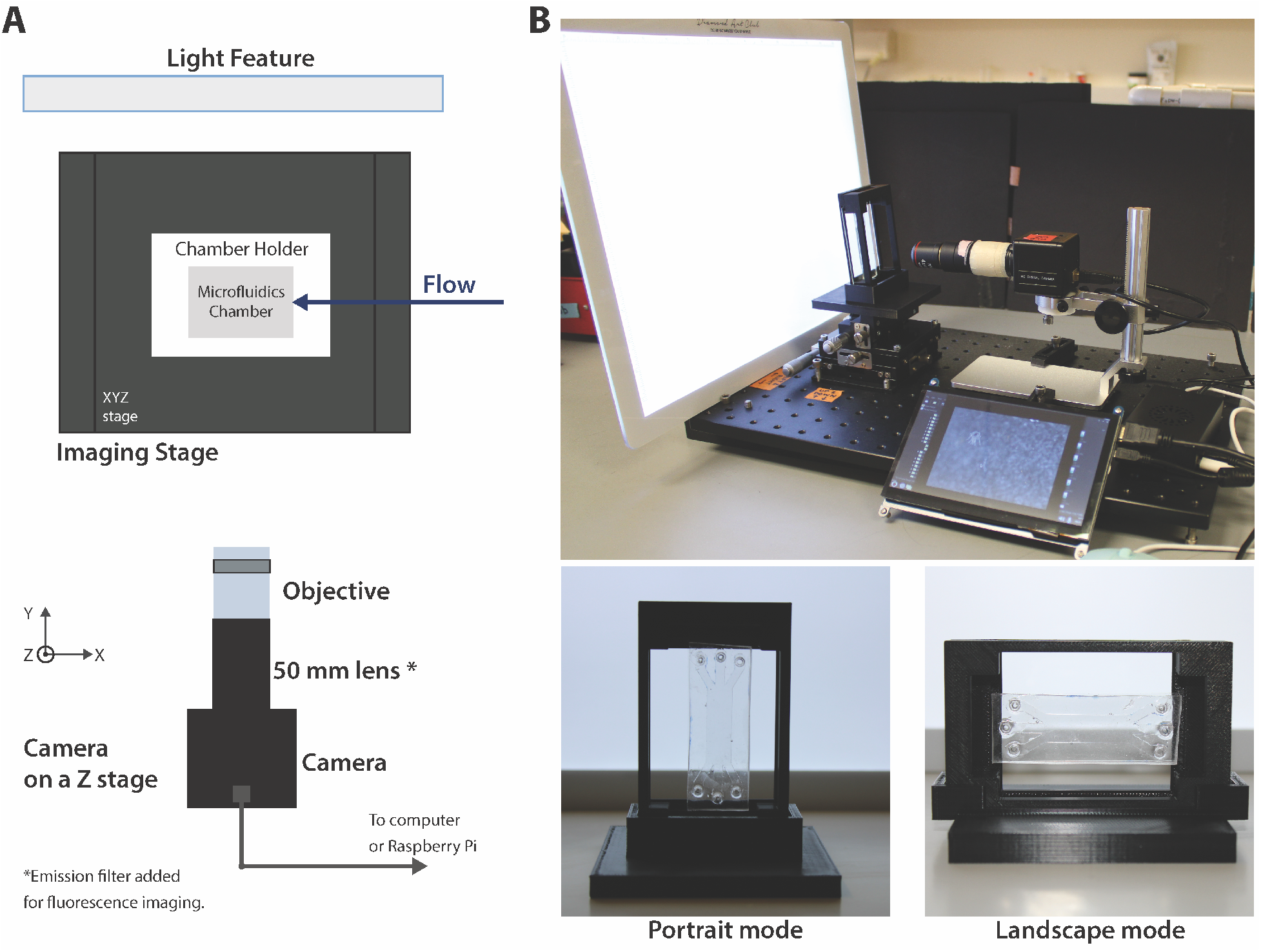
Aquavert device. (A) The system-level diagram of the Aquavert device with imaging and fluidics modules. (B) Aquavert assembled on top of an optical breadboard. The 3D-printed chamber holders were designed for the vertical positioning of standard microfluidics devices. These holders could position a chamber in either landscape or portrait mode. Aquavert enables the investigation of aquatic and marine organisms in the presence of gravity. Vertical swimming of marine organisms was of interest.

### 2.2 Imaging Module

The imaging module includes optical parts, a light source, a sensor (camera), and mechanical components for support. The imaging sensor was a webcam camera (ArduCam). Optical parts included an objective (2X, 4X, and 10X objectives were used), a tube lens (50 mm), an emission filter for fluorescence imaging (Chroma ET605/70m or Chroma ET500lp), and lens tubes for securing components. For most experiments, a light pad was used to ensure uniform illumination. An off-the-shelf LED lamp was used for fluorescence microscopy applications. A laptop (AMCap App for Windows) or Raspberry Pi was used to control the image acquisition and the camera. For the Raspberry Pi application, a simple Python code was developed. The code is available from the open-sourced project repository. Beyond cost minimization and improved portability, Raspberry Pi enabled precision control of up to two independent cameras. Supplemental Figure 1 provides images of standard USAF resolution targets for the objectives and configurations used in this investigation. The camera was attached to a Z-stage using a 3D-printed connector. Two mounting bases (Thorlabs 201/579-7227) secured the vertical stage to the optical breadboard during the experiment. The light pad was secured vertically behind the sample stage using a combination of 3D printed bases, two screws, and two Thorlabs posts. The entire system was built on an optical breadboard. Supplemental materials and the project repository provide a complete bill of materials and 3D design for all components.

### 2.3 Fluidics Module

The fluidic module includes the chamber and the mechanical support. Three primary chambers were demonstrated in the investigation. These designs were the single channel (4 mm wide, 63 mm long), two-inlet channel (4 mm wide, 63 mm long), and three-inlet channel (12 mm wide, 63 mm long). The single channel was used for baseline measurements with and without flow. The multi-inlet channels were used to create flow boundaries. The design for these chambers and alternatives are provided as supplementary materials. Fabrication protocol and the mechanical support components are described.

#### Chamber Fabrication

Laser cut (40 W CO2 desktop laser cutter) polymethyl methacrylate (PMMA; 1/16” thickness) molds were used to build chambers. Polydimethylsiloxane (PDMS, Dow Sylgard™ 184 Silicone) was cast from the PMMA molds using 10:1 mass ratio of base to curing agent. Fully cured PDMS parts were separated from the molds using a craft knife. Interfaces (inlets and outlets) were cut using biopsy punches. PDMS parts were then plasma bonded (maximum RF power 18 W) to glass slides (25 or 50-mm wide slides were used) to create the chambers. Chambers were baked at 65°C for at least 3 hours to promote adhesion between glass and PDMS further. Chambers were stored inside a Petri dish with masking tape applied to the top PDMS layer to prevent dust accumulation. Chambers were typically used multiple days after their fabrication. Silicone tubing (various inner diameters depending on the experiments) was used for all experiments. For some experiments (particularly multi-inlet chambers), 1/16” inner diameter (ID) connectors were used to prevent potential leakage at the interface between the tubing and the chamber.

#### SSMechanical support

Two different chamber holders were developed, and 3D printed to secure the microfluidics in portrait or landscape profile (Figure 1 B). Each chamber holder has a corresponding base. Before an experiment, the frame and the chamber holder were attached to the XYZ stage. The stage was used to fine-position the microfluidics chamber. We recommend that the glass slide, not the PDMS side of the chambers, face the camera during imaging. Tubing was typically attached to the chamber before the chamber was secured inside the holder. Tubing was placed behind the holder (away from the camera) to prevent obstruction or shadows. During experiments, we typically taped (using masking tape) the tubing to the breadboard to minimize vibrations or movement. For experiments with flow, the syringe pumps were placed next to (but not on top of) the optical breadboard.

Three proof-of-use experiments were developed to demonstrate the utility of the Aquavert. These experiments were (a) imaging with portrait mode chambers, (b) imaging with landscape mode chambers, and (c) fluorescence imaging all with and without flow.

### 2.4 Behaviors Inside Portrait Mode Chambers

In the first set of experiments, the chambers were positioned in portrait mode. Multiple experiments with and without flow were conducted. A summary of key demonstrative results is provided. In the first example, sand dollar and sunflower sea star larvae inside a static chamber were examined. Sand dollar larvae have an armed morphology. Arms are ciliated extensions used for swimming and feeding behaviors. Similarly, starfish larvae use ciliary bands spanning their body to swim and feed. However, unlike sand dollar larvae, they do not possess arms. Briefly, marine larvae were grown as described previously using standard protocols [21, 22]. Multiple cultures were used throughout the study. Given that the experiments were conducted at room temperature, sand dollar larvae cultures were maintained at room temperature. Low-food algal feeding treatments (1,000 cells of the alga *Rhodomonas lens* per mL) were used. The single channel had a rectangular cross-section (height 1.6 mm ×width 4 mm) and was 65 mm long. Larvae were loaded inside the channel while it was placed horizontally (flat) on the bench. The channel was partially primed with seawater then loaded with larvae. Given the dimensions, the liquid inside a vertically oriented channel would be flushed out due to gravity. We attached tubing connected to a 1 mL syringe (both filled with seawater) to the lower inlet of the channel to prevent the flushing effect. The top outlet was left disconnected and exposed to air (as apparent from the meniscus). In this example, a 2X objective was used to observe the three larvae. Figure 2 illustrates the setup and sample images from the experiment. A recording of the experiment is also provided (see Movie 1).

**Figure 2:**
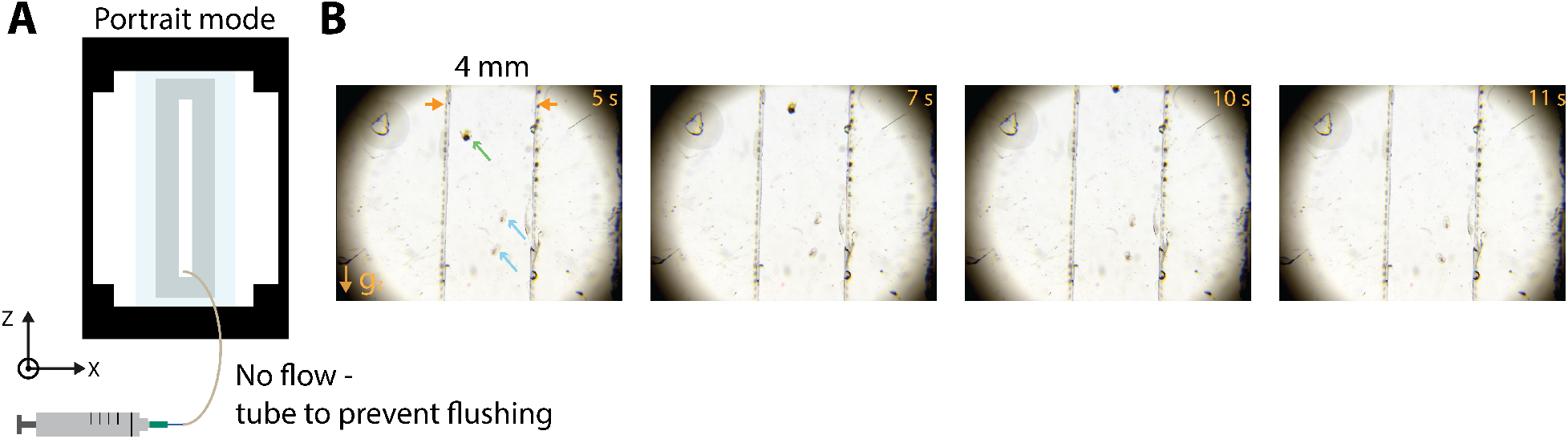
Portrait mode chambers with no flow. Vertical swimming behaviors of marine larvae without flow can be investigated via the chamber. (A) A diagram of the portrait mode chamber with no flow. The tubing was attached to prevent the chamber from flushing due to gravity. (B) A single 4 mm wide channel was used. The direction of gravity is highlighted with *g*_*z*_. The green arrow indicates a sand dollar larva. The blue arrows indicate two Sunflower sea star larvae. A 2X objective was used for imaging the chamber.

In the second example, we examined the swimming behaviors of sand dollar larvae in the presence of flow using the same single-channel design. In the ocean, larval dispersal is necessary for recruitment and metamorphosis to the juvenile stage. Before the recording, the larvae were placed inside the chamber. Next, tubing was attached to a syringe and syringe pump system. The pump moved sea water containing unicellular red algae (*Rhodomonas lens*, strain CCMP739) into the channel from the lower inlet (moving against gravity). Algae are used as the primary food to culture marine invertebrate larvae. In this case, a larva swam against the flow (sinking) with a helical path (see Movie 2). The video recording from the experiment was combined with DeepLabCut application [23] to track larval swimming (Figure 3 and see Movie 3). In the third example, we used a Y-shaped channel to create a laminar boundary. Sea water with and without algae was infused inside the chamber against gravity (bottom to top) at a rate of 0.25 mL/hr (Figure 4A provides the diagram). Algae here also enabled boundary visualizations. In this example, the larva’s initial position was on the side containing algae. The larva swam across the boundary to the side without algae (Figure 4B). In other experiments, the opposite behavior was observed. We plan to use Aquavert to characterize these behaviors in the future. These experiments confirmed the feasibility of using vertical channels in portrait orientation.

**Figure 3:**
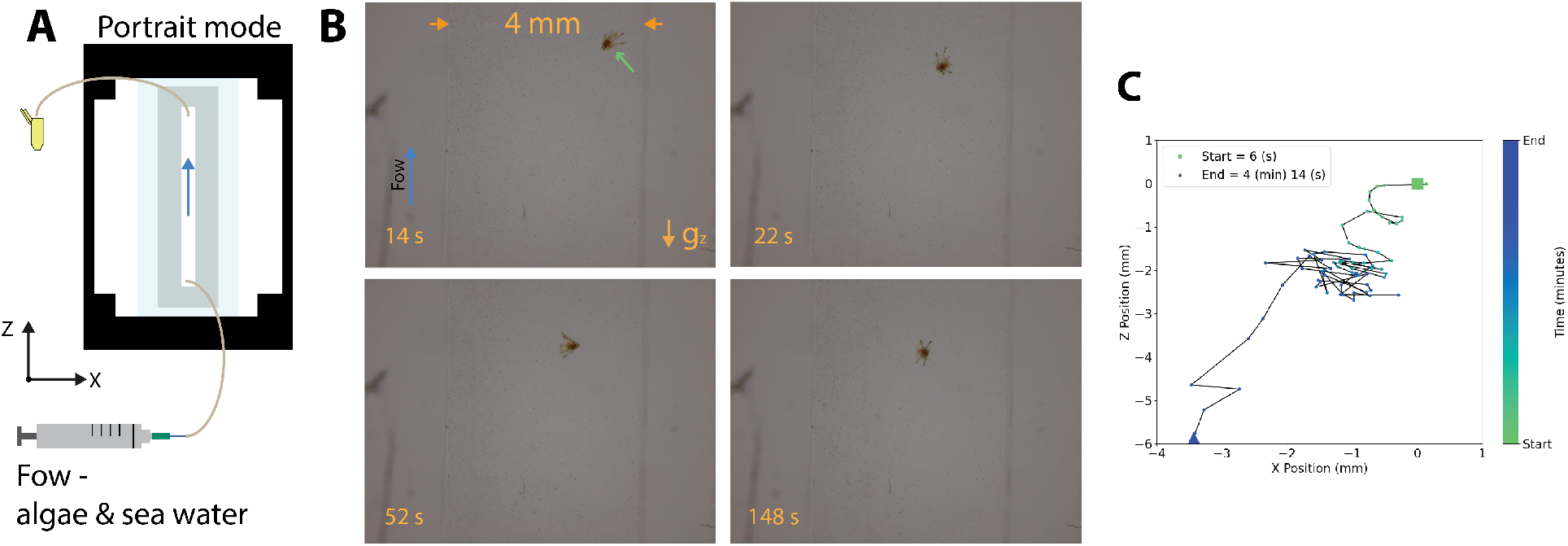
Portrait mode chambers with flow. Vertical swimming behaviors of a sand dollar larva in the presence of flow. (A) A diagram of the portrait mode chamber with flow. Sea water containing red algae was continuously infused inside the chamber. The blue arrow indicates the direction of the flow against the gravity. The direction of gravity is highlighted with *g*_*z*_.(B) A 4 mm wide channel was used (orange horizontal arrows). In this example, sand dollar larvae primarily swam across the channel’s height in the Z-direction against the direction of flow in a helical path. (C) The position of the larva inside the channel was tracked for the duration of the video. The trajectory path line demonstrated the complex swimming pattern of the organism. A 4X objective was used for imaging the chamber.

**Figure 4:**
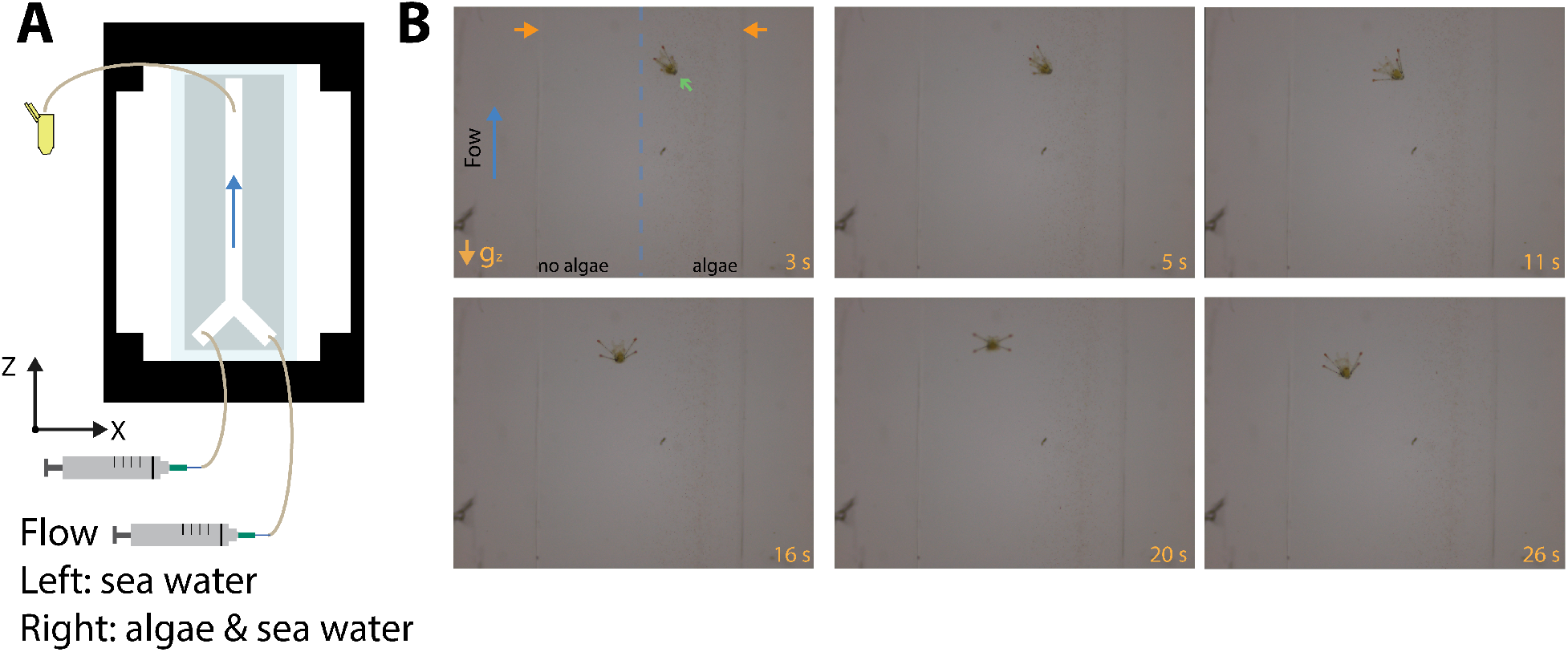
Portrait mode chambers with flow and boundaries. (A) A Y-shaped device was used to create a boundary and regions with and without algae. Sea water containing red algae were continuously infused inside the chamber at 0.25 mL/hr. The blue arrow indicates the direction of the flow against the gravity. The direction of gravity is highlighted with *g*_*z*_. The segments with and without algae are highlighted with a dashed line. The chamber is 4 mm wide (orange horizontal arrows). (B) In this example, the sand dollar larva primarily swam across the boundary from region with algae to the region without algae.

### 2.5 Behaviors Inside Landscape Mode Chambers

In the second set of experiments, the chambers were positioned in the landscape mode. Investigating the interactions between vertically swimming microorganisms with horizontal shear flows is of great interest [16, 24, 25]. Fluid flow in this profile could establish vertical gradients in horizontal velocity (i.e.,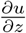). Aquavert enables precision control over these horizontal shear flows and corresponding vertical gradients.

First, a single channel (4 mm wide, 65 mm long) was used. Similar to portrait mode experiments, the larvae were loaded before the channel was vertically positioned. Sea water containing algae was continuously infused inside the chamber. Figure 5 (A and B) provides these results. To prevent accidental flushing due to gravity, tubing was attached to inlets and outlets, even for experiments without flow.

**Figure 5:**
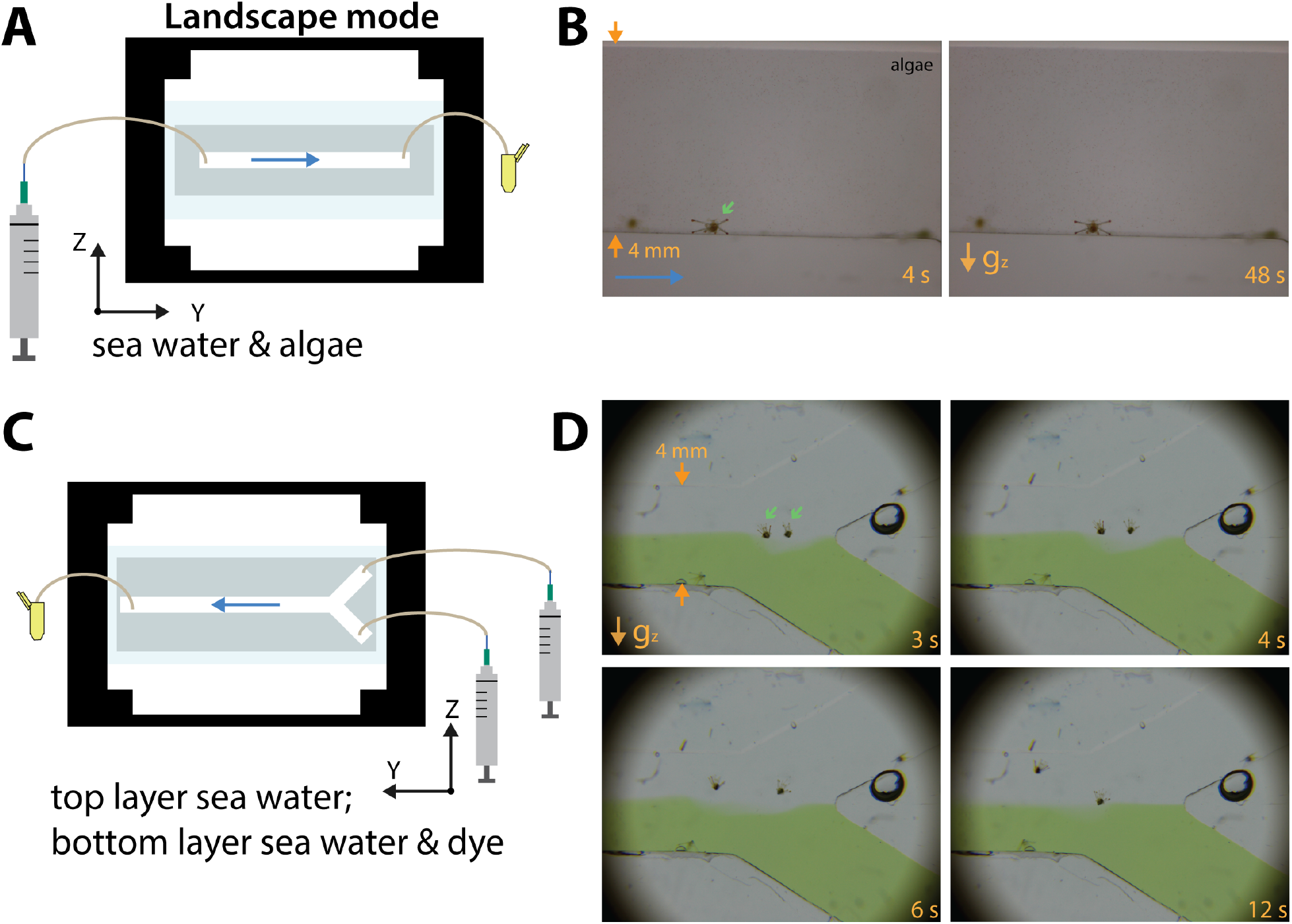
Landscape mode chambers. (A) The diagram of landscape mode single channel with flow. (B) Sea water containing algae was continuously infused across the channel. The sand dollar larva inside the channel is indicated with a green arrow. (C) A Y-shaped device was used to create a boundary and regions with and without indicator dye. (D) In this example, the sand dollar larva primarily swam across the boundary from the region with algae to the region without.

Microfluidic chambers are dominated by laminar flows, which enable the introduction of well-controlled flow profiles, fluid augmentations, and predictable flow boundaries. These profiles and well-established boundaries are valuable for investigating swimming and modulating the content of each fluid segment (food or soluble factors). As part of the Aquavert device, multiple chambers (a Y-shaped two-inlet device and a three-inlet trident-shaped device) were developed to create fluid segmentation. Key results are provided in Figure 5 (C and D). In this case, trace amounts of food dye were added to the bottom layer to enable boundary visualizations. The volumetric flow rates in the top and bottom layers were 0.5 mL/hr and 1.0 mL/hr respectively. We experimentally observed a stable boundary. Additionally, Reynolds number (Re) estimations suggested that the maximum Re inside the chamber was less than 1, corresponding to laminar flow. Three sand dollar larvae were introduced from the lower inlet in this experiment. As demonstrated in Figure 5 D, two larvae navigated across the boundary from the bottom to the top layer (see Movie 4). The flow in this experiment was directed parallel to the Y-direction (i.e., horizontal stratification).

To further demonstrate the device application, the behavior of freshwater *Hydra* inside the 4 mm wide channels (landscape orientation) was examined (see Supplementary figures). While not a marine organism, results from *Hydra* demonstrated that the device can be used with other organisms, emphasizing the role of gravity, which is often ignored in conventional plate or microfluidic experiments.

### 2.6 Fluorescence microscopy

Beyond tracking the organisms, visualizing the flow patterns they generate as they eat or swim can provide key functional information on their behaviors. To this end, PIV systems, conventional fluorescence microscopes, and DIY systems have been used[14, 26, 27, 28]. We used the modularity of the Aquavert to enable fluorescence imaging. To this aim, first, we replaced the light source with an off-the-shelf external LED lamp for illumination. The lamp was selected because it provided various color selections and brightness adjustments. Alternative light sources are available, and recommendations are provided in the bill of materials. Second, we added an emission filter as part of the detection path. The choice of the filter was informed by the red fluorescent beads (542 nm ex./ 612 nm em.; 1.1, 2, and 15 *µ*m diameters; Fluoro-Max) that were used in this investigation. Given the beads’ reported excitation wavelength, the LED was set to green. Key results are presented to demonstrate the device’s capabilities. Here, the use of Raspberry Pi was valuable for precision control of acquisition details.

First, a micro-fluorescence cuvette (Thor labs CV10Q7FA) was used to validate the feasibility of the fluorescent imaging using the LED lamp. A sand dollar larva was added to the cuvette containing beads and sea water. A sample recording from these experiments and the corresponding pathline particle tracking is provided which includes qualitative visualization using the Flowtrace algorithm [29] (Movie 5). We used single or Y channel devices in landscape mode. Figure 6 provides sample results with and without flow. Here data were collected both via a laptop (AMCap App for Windows) or a Raspberry Pi using our Python program using the OpenCV library. The Raspberry Pi system was valuable since it provided fine control over the acquisition details. In one example, beads were introduced from one of the inlets of a Y-shaped device (Movie 6). Results from this experiment were combined with a PIV analysis approach [30] to better visualize the flow of the beads inside the channel.

**Figure 6:**
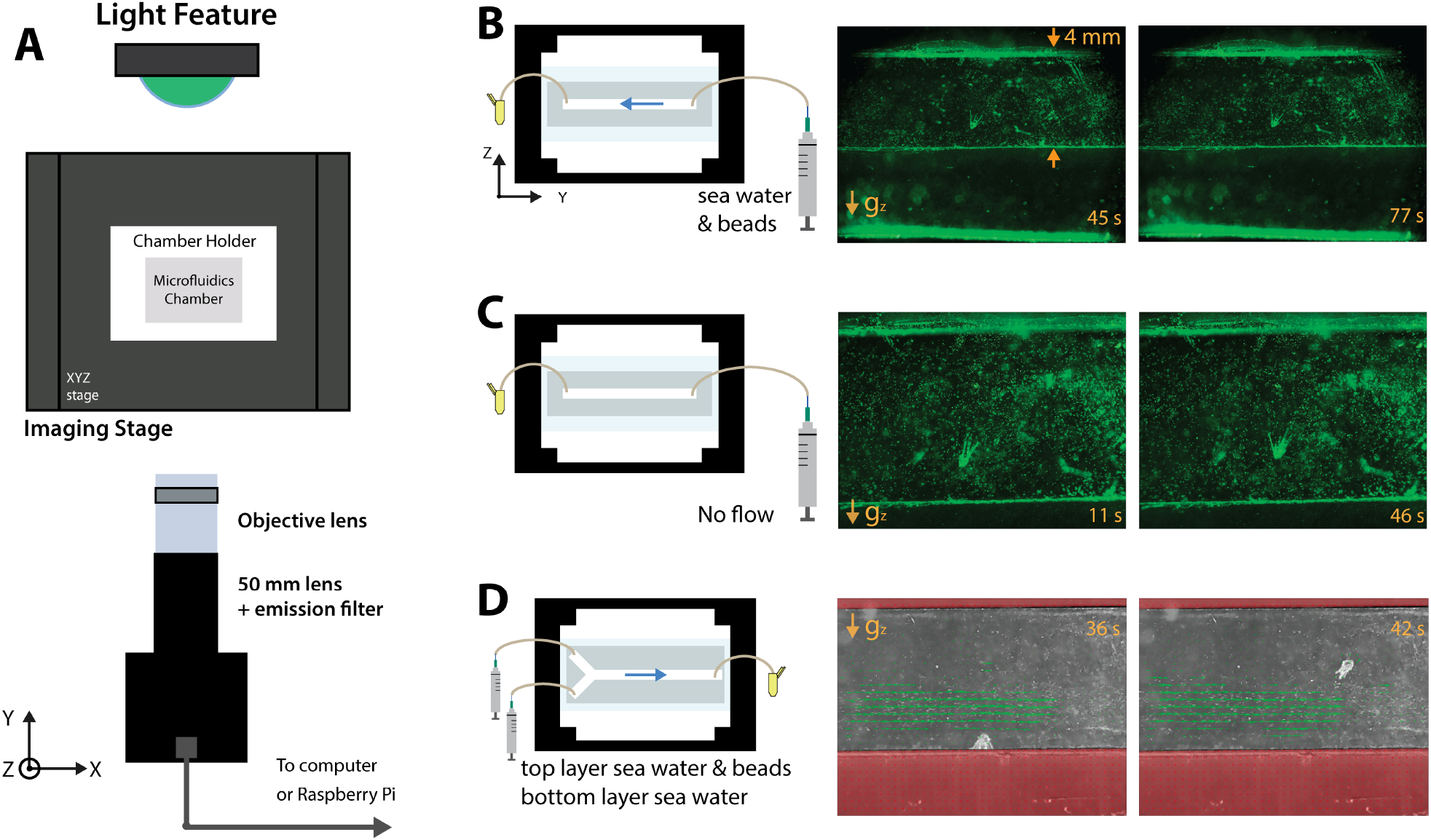
Fluorescence imaging. (A) The updated system-level diagram for fluorescence imaging. (B) An example of a landscape mode chamber with flow with beads and seawater. The flow rate was at 5 mL/hr. The direction of gravity is highlighted with *g*_*z*_. The white arrow indicates the sand dollar larva.The channel width (4 mm) is highlighted. A 2Xobjective lens was used. (C) An example of a landscape mode chamber with no flow. A 4X objective lens was used. (D)An example of PIV analysis for a segmented channel (with and without beads).

## 3 Discussions and conclusion

We developed a device (Aquavert) that combines DIY imaging and microfluidics to study marine organisms’ behaviors, emphasizing vertical swimming and interactions with fl under well-controlled conditions. Our efforts were inspired by the circular chamber developed by Krishnamurthy and co-authors [17, 18]. We used standard microfluidics instead of the ring-shaped chamber developed by Krishnamurthy to control flow profiles and corresponding boundaries. Unlike conventional microfluidics experiments, the chambers were secured vertically to enable vertical swimming. Chambers could be positioned in either landscape or portrait format to introduce flow profiles parallel to the x-axis or z-axis. While the chambers limited the total path an organism could swim, it enabled the introduction of external flows and their corresponding segments. Fluid segments with and without food (algae) were created using the laminar flow profiles. These boundaries provide an approach to investigating feeding and swimming behaviors under well-controlled conditions. Investigating the feeding behaviors of marine larvae is critical for several biological inquiries (e.g., functional morphology and population ecology) [31]. We prioritized modularity as our key goal. The rationale behind this decision was to enable easy switching of components. Additionally, control with a Raspberry Pi was valuable for cost, size reduction, and potential control of multiple cameras.

There are limitations in the current version of the device presented in the manuscript. First, automated tracking similar to existing efforts [17, 6, 4] could be a valuable addition. An alternative strategy is to use multiple detection arms to visualize the entire chamber. This strategy is feasible given the low cost of the components and simultaneous acquisition control that can be implemented with a Raspberry Pi. One concern related to moving parts is the miscalibration or misalignment of parts over time. Second, a mesh similar to the Meioflume device [32] could be introduced at inlets and outlets to prevent the organisms from being washed out of a chamber. Alternatively, the fluid inside the chamber and the organism can be recirculated. Third, temperature control strategies could be included in the systems. In this investigation, we used sand dollar larvae as a model organism since they can be cultured and handled at room temperature. However, other organisms (e.g., sea urchin larvae) are more sensitive to temperature variations and are typically investigated in 16°C. In one low-cost strategy, a Peltier cooler can be used to maintain the temperature of the liquid reservoirs. The ability to manipulate temperature, either sequentially or at the same time, in different flow segments is of great value since it can enable investigations related to temperature preference or the role of viscosity in feeding. In another approach, the entire system can be operated inside a cold room. Additionally, the device can be modified to explore the effects of temperature gradients across an observation chamber. For example, heat sink and source channels parallel to the observation chamber can be implemented to such gradients or maintain a desirable temperature. Finally, the total volume and size of the chambers can be increased. In the context of marine larvae, confinement is often a concern. We used chambers that were much larger than typical microfluidics. Specifically, the smallest dimension (chamber height) was about 10x bigger than typical larvae (estimating a larva length scale at 200 *µ*m). However, carefully considering a chamber size and the aim of an investigation is important. Our goal for the current experiment was to use microscopy to visualize a larva fully. To address some experimental questions only the position of a larva may be needed. In such cases, the chamber dimensions can be extended.

There is a broad interest in fluid dynamics (swimming and feeding) of microorganisms ranging from single-celled organisms, zooplankton, and marine larvae [33, 34, 35]. Lab-based investigations, with all their limitations, provide pathways to study these organisms under well-controlled conditions. Aquavert combines the effects of flow and gravity. The introduction of the well-controlled flow provides stable segments and neighboring regions within the chamber with and without factors ranging from temperature, flow forces, food, microplastics, or other chemicals. Investigating the interaction of marine microorganisms with these boundaries in the presence of gravity (i.e., gravitactic behaviors) can provide critical insights about organism behaviors in their natural environments. Experimental results could also be used to examine against previously developed models (gravity-buoyancy and drag-gravity models) [36]. The approach presented, particularly devices in landscape mode with flow, could be used to investigate the formation of phytoplankton thin layers at the microscale. In this context, the combination of microscopy and well-controlled flow could provide high-resolution imaging of phytoplankton to enable examination of behaviors (e.g., gyrotactic trapping or convergence of swimming) related to thin layer formation [37]. Beyond marine biology applications, the device could enable other microfluidic investigations where the role of gravity is essential (e.g., gas bubbles or plant root formation[38]).

## Acknowledgments

Authors express gratitude to members of Ahrar-lab (Sandra Trajkovski and Emily Chrasta) and members of Pace-lab (Ivanna Arrizon Elizarraras). Authors are thankful to CSULB Marine Lab.

Authors express gratitude to Prof. Bruno Pernet and Prof. Rob Steele for their support and feedback on the manuscript.

This work was supported in part by a CSUPERB new investigator grant to S. Ahrar; a CSULB COAST student award to H. Obenshain and B. Lopez; and a STEM-NET collaborative seed award from the CSU Office of the Chancellor to S. Ahrar and D. Pace.

## OSF Repository

OSF Name: Link: to be added.

As part of the project, resources, including the complete library of videos, design files, and code for analysis, have been shared.

## Conflict of interest

Authors declare no conflict of interest.

## Institutional Review Board Statement

Not applicable.

## List of Supplementary Movies

- Movie 1: Marine larvae inside static chamber.
- Movie 2: A sand dollar larva inside a portrait mode chamber.
- Movie 3: A sand dollar larva inside a portrait mode flow-segmented chamber.
- Movie 4: Sand dollar larvae inside flow-segmented landscape mode chamber.
- Movie 5: Validating fluorescence imaging - larva inside a cuvette.
- Movie 6: Fluorescence imaging inside a landscape mode device.

## List of Electronic Supplementary Information (ESI)

- ESI 1: Chamber designs.
- ESI 2: 3D printed components.
- ESI 3: Bill of materials.

---

**Figure 7: Sup Figure 1:**
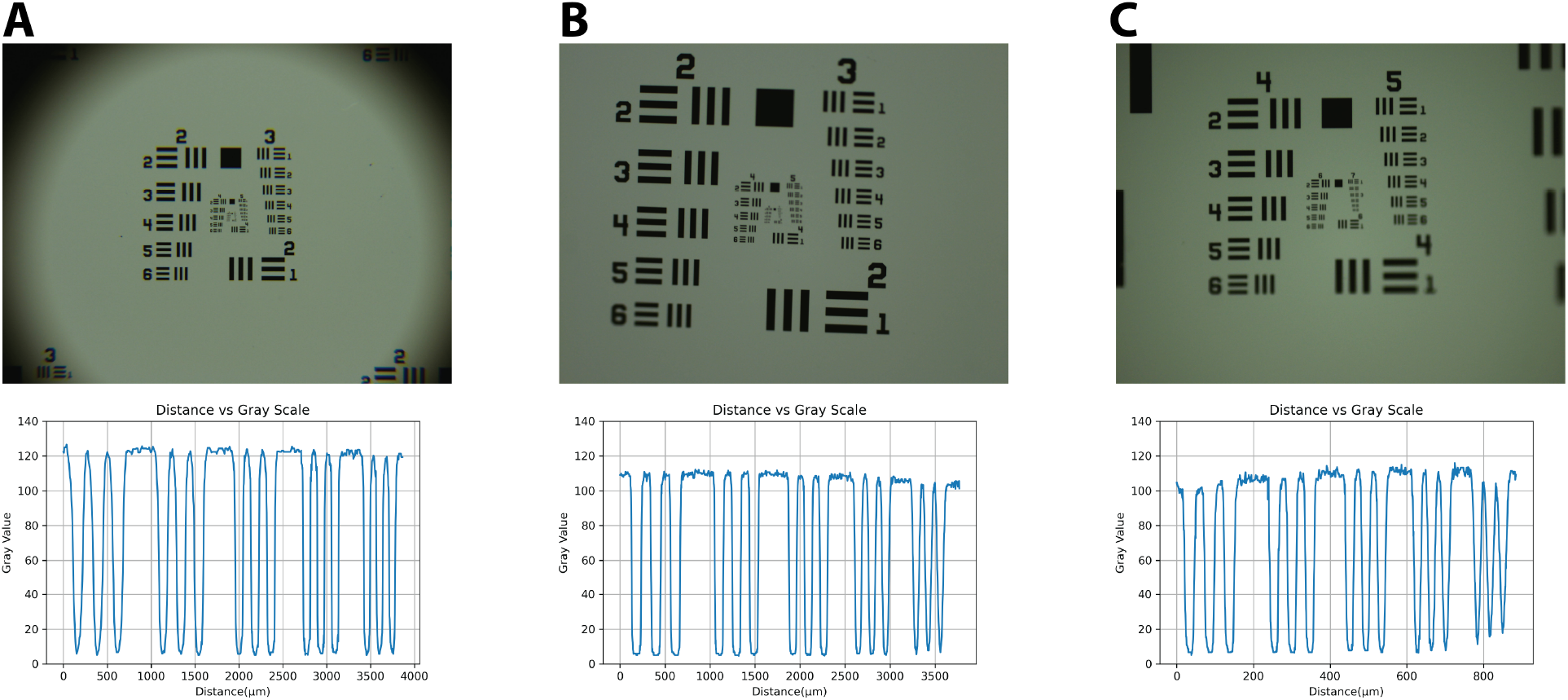
Images of USAF resolution slide. Sample images were obtained via the objectives that were used throughout the manuscript. Given the modularity of the system, both objectives and the tube lens can be readily adjusted. (A) Images via a 2X objective. (B) Images via a 4X objective. (C) Images via a 10X objective.

**Figure 8: Sup Figure 2:**
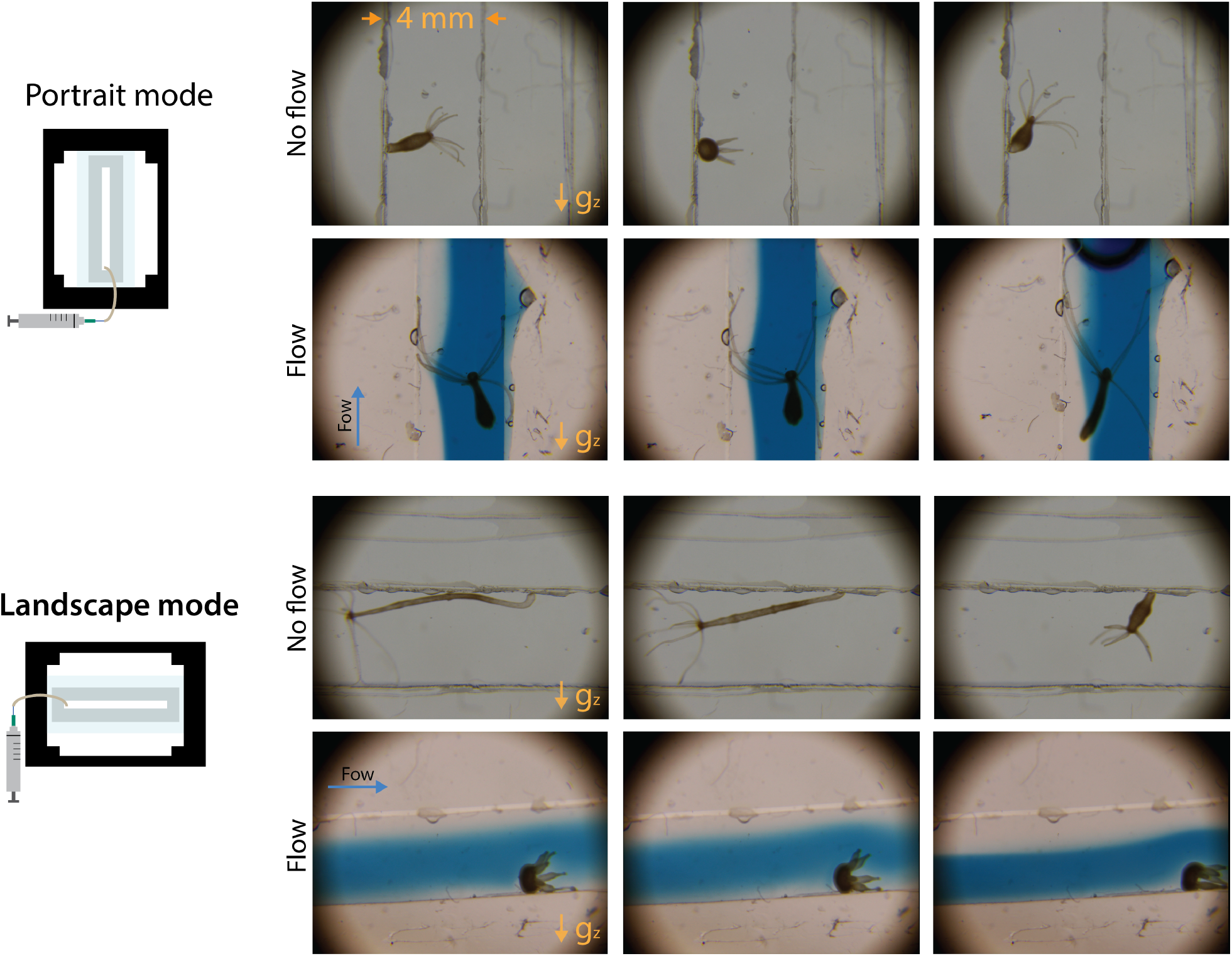
Imaging *Hydra* with Aquavert. Organisms were visualized inside 4 mm wide chambers with and without flow. Both portrait and landscape orientation were validated. Results extended the application of Aquavert beyond marine larvae.

## Notes

### Competing Interest Statement

The authors have declared no competing interest.

